# fMRI hyperscanning reveals that coordinated exploration predicts successful conversation

**DOI:** 10.64898/2026.11.18.767620

**Authors:** Sebastian P.H. Speer, Caroline Lee, Laetitia Mwilambwe-Tshilobo, Lily Tsoi, Shannon M. Burns, Emily B. Falk, Chris Baldassano, Diana I. Tamir

## Abstract

Conversations among friends are often more dynamic, enjoyable, and wide-ranging than those between strangers. How do friends do this? We propose that inside jokes, shared experiences, and mutual interests give friends an aligned mental map, allowing them to leap between topics without losing each other. Strangers may begin in separate mental landscapes, limiting opportunities for exploration as they take smaller steps to avoid confusion. To test these possibilities, we tracked the dyadic dynamics of friends (n = 60) and strangers (n = 54) as they engaged in a real-time conversation using fMRI hyperscanning. We employed Hyper-Hidden-Markov-Modeling to project partner’s neural states into a shared latent space and to segment them into discrete events. This allowed us to quantify how similarly each dyad represented each event (representational alignment), and how spread out conversational events were in latent space (representational exploration). We tested how neural alignment predicted exploration, how both neural measures relate to the breadth of topics discussed, and how they predicted the subjective experience of the conversation quality in friends and strangers. We found that the more a dyad aligned neurally, the more they also explored. While both neural alignment and exploration predicted the breadth of conversations, only neural exploration predicted better perceived conversation quality. Friends were more neurally aligned, which was associated with more exploration. Thus, when friends’ brains represent the world more similarly to each other, they tend to embark on more diverse and far-reaching conversations spanning a broader range of topics, all while staying anchored in common ground.

**Significance Statement:** Good social interactions create social bonds that help people live longer with less stress and lower rates of disease. Yet, how conversations bring people together remains unclear. Previous research has proposed two seemingly opposing routes: Alignment and exploration. Scanning conversation partners’ brains simultaneously, we found that partners with more aligned neural representations explored a wider range of neural states. Greater exploration but not alignment was associated with stronger social connection. Together, these findings suggest that alignment and exploration are complementary, supporting an account of coordinated exploration in which partners venture into new territory together while remaining anchored in common ground. This work clarifies the neural dynamics of successful conversation and could inform strategies to strengthen social bonds and reduce loneliness.

## Introduction

From trying to charm a first date to catching up with a grandparent, the quality of a conversation can shape the connection that emerges from it. Good conversations bring people joy, strengthen relationships, support mental and physical health, and help hold groups together (Baumeister et al., 2007; Diener & Seligman, 2002; Leary & Baumeister, 2000; Ryff, 1995; Sandstrom & Dunn, 2014; Yu, 2023). But what makes a conversation good?

### From alignment and exploration to coordinated exploration

One prominent answer put forward by prior research is that good conversations depend on alignment between conversation partners. Alignment refers to the way people spontaneously synchronize their body movements (Ramseyer & Tschacher, 2011; Tunçgenç & Cohen, 2018), language (Doré & Morris, 2018; Manson et al., 2013), physiology (Behrens et al., 2020; Chen et al., 2021; Prochazkova et al., 2021), and neural responses (Kinreich et al., 2017; Liu et al., 2017; Moffat et al., 2026; Wang et al., 2022; Zhang et al., 2018). Alignment is thought to make conversations more fluent (daSilva & Wood, 2024; Koban et al., 2019), reduce the cognitive effort required to understand the partner (Brown-Schmidt et al., 2015; daSilva & Wood, 2024; Garrod & Pickering, 2004; Templeton et al., 2022), and increase partners’ perceived similarity (Chartrand & Bargh, 1999; Cornejo et al., 2018; daSilva & Wood, 2024). These effects strengthen social connection. Alignment has been linked to emotional support, interpersonal liking, social influence, and social cohesion (Doré & Morris, 2018; Hoehl et al., 2021; Ireland et al., 2011; Ludwig et al., 2013; Prochazkova et al., 2021; Street et al., 1983). Alignment, therefore, appears to be a promising route to successful conversations.

That said, if two conversation partners repeated and validated everything their partner said, they would be highly synchronized, but the conversation would quickly become predictable and boring. Indeed, recent research has highlighted exploration as an additional route to good conversations (Ravreby et al., 2022; Sened et al., 2024; Speer et al., 2024, 2026). Exploration refers to the ways conversation partners push and pull one another in new directions, for example, by introducing new ideas, considering distinct viewpoints, and avoiding repetition (Burns et al., 2025; Speer et al., 2024, 2025, 2026). Exploration may enhance conversation quality by making interactions more engaging and surprising and by creating opportunities for partners to learn about one another (Poli et al., 2024; Speer et al., 2024, 2025; Templeton et al., 2024; Tsang et al., 2025). Exploration may also improve emotion regulation when one partner introduces emotions or perspectives that counterbalance the other’s (D. S. Lee & Fujita, 2023; Powers & LaBar, 2019). If one person is sad, their partner can console them by talking about more positive topics and thus pulling them into brighter emotional terrain, rather than co-ruminating in a way that entrenches the sadness (DiGiovanni et al., 2024, 2026). Indeed, neural and topic exploration in conversation has been associated with better social outcomes such as increased liking, enjoyment (Sened et al., 2024; Speer et al., 2024), and agreement (Speer et al., 2026). Together, this evidence points to exploration as an important additional route to more successful conversations.

These two accounts suggest distinct, but not necessarily incompatible, routes to social connection (Kinreich et al., 2017; Parkinson et al., 2018a; Ravreby et al., 2022; Sened et al., 2024; Shen et al., 2025; Sievers et al., 2024; Speer et al., 2024, 2025, 2026; Stephens et al., 2010). Yet, because these two processes have largely been studied in separate literatures, prior work has not yet established how they might coexist within the same conversation. As a result, central questions remain unresolved: in what ways do these two processes relate to one another (if at all), and how do they relate to social connection? At least three possibilities exist. First, alignment and exploration may operate largely independently, perhaps at different spatial levels of neural representation (e.g., voxels patterns, regions, or networks; Song et al., 2025) or over different timescales. Second, they may compete: exploration may disrupt alignment when partners depart from shared states and move in different directions. Third, alignment may facilitate exploration: a shared representation of the current conversational event may allow partners to move together into new conceptual, emotional, or neural states without losing mutual orientation. We refer to this pattern of movement into new territory while partners remain mutually oriented as *coordinated exploration.* We favor the third possibility, proposing that alignment and exploration function as complementary dynamics whose coordination may foster social connection.

### Friendship as a context for coordinated exploration

We leverage variation in people’s social history as a valuable context for evaluating this account. Specifically, friendship may modulate the extent to which the two dynamics are engaged in conversation. Friends enter conversations with a richer base of shared experiences, references, and expectations, embedded in a distinctive micro-culture of jointly created meanings (Rossignac-Milon et al., 2026). Strangers, in contrast, must establish common ground with an unfamiliar partner during the interaction. Having more baseline common ground may allow friends to represent ongoing conversational content more similarly from the get-go, and to explore new conceptual territory without losing mutual orientation. Consistent with this account, friends show greater similarity in their neural time course during passive exposure to naturalistic stimuli (Parkinson et al., 2018b; Shen et al., 2025) and greater neural representational alignment at the beginning of conversations (Speer et al., 2024). At the same time, friends *also* explore more distinct neural space than strangers (Speer et al., 2024). In the current study, we used fMRI hyperscanning to track the neural dynamics of dyads of friends and strangers as they engaged in naturalistic, real-time conversation. We predicted that friends would exhibit *both* heightened neural alignment and neural exploration, relative to strangers.

### Current methodological challenges

Testing these predictions requires a framework that captures the temporal structure of conversation, localizes its neural dynamics, and quantifies alignment and exploration simultaneously as a conversation unfolds.

First, measuring conversational dynamics requires identifying meaningful conversation events for each partner. Most decoding approaches are not sensitive to the natural structure of conversation: they are applied either to each TR separately or to pre-defined events, without estimating how segments are neurally aligned across people. To address this, we must have a way to define conversational events that map onto naturally flowing conversation. Here, we use the term conversational event to refer to a temporally bounded segment of the conversation in which partners are engaged with a coherent idea, memory, topic, or perspective. These events do not correspond to independent imaging volumes (TRs), nor do they necessarily map cleanly onto experimenter-defined windows such as turns or topics. Instead, each partner may enter, maintain, and leave the same conversational event at different times.

Second, understanding the mechanistic biological link between conversation dynamics and social connection requires localizing these dynamics to specific neural systems. Whole-brain measures can characterize these dynamics but may obscure contributions of individual regions or networks. By identifying the neural systems in which alignment and exploration operate, we gain insight into the potential psychological processes that support them, such as semantic representation, mentalizing, attention reorientation, control, affect, or sensorimotor processing, among others.

Third, testing how alignment and exploration relate to one another requires measuring both within the same representational framework. Existing neuroimaging approaches, however, have generally examined the two processes separately.

Alignment between conversation partners is typically studied using localized *intersubject correlation* (ISC; Nguyen et al., 2021; Parkinson et al., 2018b; Shaw et al., 2018; Shen et al., 2025; Sievers et al., 2024; Xie et al., 2020), which measures mutual engagement by correlating activity in corresponding brain regions across two people. *Linguistic encoding models* likewise capture the extent to which language aligns partners’ neural responses (Zada et al., 2024, 2026). These approaches are agnostic to exploration because they quantify whether partners’ neural responses are coupled to one another, or to the linguistic content of the conversation, but not how each partner’s neural state moves through representational space. As a result, they cannot capture how the two processes co-occur or interact.

*Mental state decoding* (Speer et al., 2024, 2026) addresses this limitation by translating participants’ whole-brain neural activity into coordinates in a psychologically interpretable three-dimensional space (e.g., mental state space; Tamir et al., 2016; Tamir & Thornton, 2018; Thornton et al., 2022). As a result, this approach enables quantifying alignment and exploration within the same representational framework. It therefore demonstrates how both dynamics can be measured together, but leaves unresolved their neural localization and the correspondence of conversational events across partners. Testing our predictions therefore requires a framework that combines this joint quantification with neural localization and sensitivity to the temporal structure of conversation.

### A shared framework for measuring neural alignment and exploration

The current study employed a novel computational approach termed Hyper-Hidden-Markov Modeling (HHMM, Lee et al., 2023), which offers three key advantages over prior methods. First, it is sensitive to the temporal structure of conversation. HHMM segments neural timecourses into discrete event states and aligns corresponding states across conversation partners, allowing us to compare neural patterns associated with shared conversational events rather than isolated TRs or externally imposed windows (see Figure 1). Second, HHMM localizes these dynamics to specific neural systems. We conducted our analysis at the network level using Yeo’s seven-network parcellation (Yeo et al., 2011), because our hypotheses concerned broad functional systems supporting conversation rather than fine-grained regional differences. This network-level approach also provides more stable event representations by pooling across functionally related regions, reducing parcel-level noise, and mitigating the multiple-comparisons burden. Third, and most importantly, HHMM allows us to quantify alignment and exploration directly within the same representational framework. After identifying shared event states, HHMM projects the neural patterns associated with those events into a shared latent space, where we can quantify both alignment and exploration. Alignment captures how similarly conversation partners represent what they are currently discussing. It is measured at the level of individual conversation events, with greater alignment reflected in partners’ neural event patterns occupying closer locations in the shared latent space (for details see Methods and Results). Exploration, instead, captures the breadth of neural representation elicited across the conversion as a whole. It is measured as the total area of the shared latent space covered by the dyad’s neural event patterns (for details, see Methods and Results). This joint quantification is what allows us to directly test our central hypothesis that alignment and exploration are complementary conversational dynamics. Further, we validate these latent neural measures with an observable, established feature of conversation: the extent to which partners explore different topics (Speer et al., 2024, 2026).

**Figure 1.**
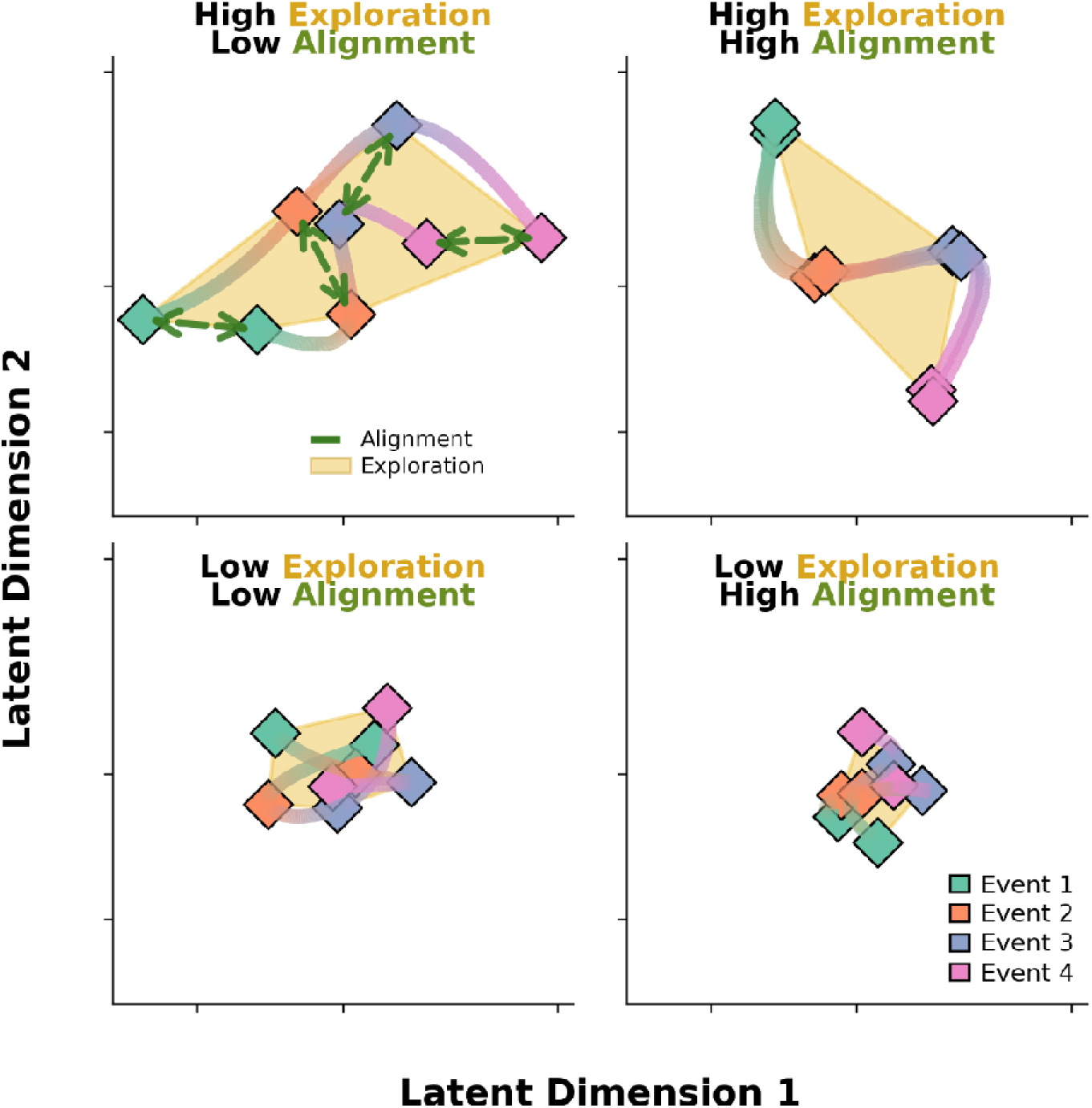
HHMM projects events into a shared representational space. In this space, we measure neural alignment as the proximity between corresponding events (green dashed arrows). In the same space, we also measure neural exploration as the total area covered by all the events in a conversation (golden area). The top left panel represents high exploration and low alignment. The top right panel shows high exploration and high alignment. The bottom left panel represents low exploration and low alignment. The bottom right panel represents low exploration and high alignment.

In this study, we combined fMRI hyperscanning with HHMM to measure both partners’ neural activity during the same live conversation and estimate event-level neural trajectories within localized functional networks. We recruited 30 friends and 27 stranger dyads and presented them with prompts from the Fast Friends procedure, designed to promote interpersonal closeness (Aron et al., 1997). Using this approach we addressed four questions. First, we tested whether neural alignment and neural exploration are inversely related or instead positively coupled, which would suggest that they work together in complementary ways. Second, we examined whether neural alignment and neural exploration were associated with topic exploration during the conversation, providing a cross-level test of convergent validity. Third, we tested whether neural alignment and exploration predicted people’s assessments of conversation quality after the conversation ended. In the context of the broader hyperscanning literature (Kinreich et al., 2017; Moffat et al., 2026; Nguyen et al., 2021; Parkinson et al., 2018b; Shen et al., 2025; Sievers et al., 2024; Xie et al., 2020) and recent fMRI hyperscanning studies (Speer et al., 2024, 2026), we hypothesize that both alignment and exploration lead to higher conversation quality. Finally, we examined whether a shared social history was associated with more alignment and exploration by comparing conversations between friends to conversations between strangers.

## Results

### Dyads with higher neural alignment explore more shared latent space

Our first analysis tested the relation between neural alignment and neural exploration. We quantified alignment as the distance in shared latent space between partners’ representations of the same conversational event, such that smaller distances indicate more similar representations of what is currently being discussed (see Figure 1). For instance, if two friends are reminiscing about Italian cuisine during a shared trip to Rome, this conversational event may activate the same landmarks in their shared mental map: the restaurant they visited, the jokes they made, and their emotions linked to that experience. Their neural representations of the event should therefore occupy nearby locations in the shared latent space. By contrast, two strangers discussing Italian cuisine may share general topic knowledge, but this conversation segment is likely to evoke more idiosyncratic associations in each person, such as different restaurants, travel memories, films, or cultural stereotypes. Their representations should therefore occupy more distant locations in the latent space. We refer to this event-level similarity between partners’ neural representations as neural *alignment*.

We quantified exploration as the overall shared latent space the dyads traversed (see Figure 1). We computed the combined area of shared representational space covered by all of the dyad’s conversation events. Continuing the example, both friends and strangers might begin with Italian cuisine. For friends, however, a memory of a meal in Rome could activate distant nodes in their shared mental map: a trip they took together, the people they met there, the clothes they wore, and more global fashion trends. Because they share a mental map, friends can move from the initial topic into more distant regions more quickly. By contrast, strangers must rely more on immediately available topic knowledge and may therefore take smaller conceptual steps from Italian cuisine to Italian wine and then to the weather in Italy’s wine-producing regions, covering a narrower region of shared latent space. Neurally, higher exploration would correspond to more dissimilar activation patterns (i.e., different subsets of voxels (de)activated) from one conversational event to the next within a network. Conversely, lower exploration would correspond to neural patterns that remain more similar across events. We term this measure *neural exploration*. Together, these measures provide a framework for testing how alignment and exploration jointly unfold during conversation and how each relates to conversation.

If neural alignment provides partners with a shared mental map, then more aligned dyads should be able to move into new regions of representational space while remaining mutually oriented, which we refer to as *coordinated exploration*. If, instead, they reflect competing or independent dynamics, they should be negatively correlated or not correlated, respectively. To arbitrate between these possibilities, we used a multilevel modeling approach (see Methods for details). We first conducted a model comparison to determine whether we would treat all ROIs together, or allow them to have differing slopes. This comparison revealed that the relationship between alignment and exploration differed significantly across ROIs, χ²(6) = 174.72, p < .001, so the interaction model, where a separate slope for each network was fit, was therefore retained. Alignment positively predicted exploration in the Default (β_std_ = 0.29, SE = 0.02, 95% CI [0.25, 0.33], p < 0.001), Dorsal Attention (β_std_ = 0.16, SE = 0.02, 95% CI [0.11, 0.21], p < 0.001), Frontoparietal Control (β_std_ = 0.14, SE = 0.02, 95% CI [0.10, 0.19], p < 0.001), Somatomotor (β_std_ = 0.14, SE = 0.02, 95% CI [0.10, 0.19], p < 0.001), Ventral Attention (β_std_ = 0.10, SE = 0.02, 95% CI [0.05, 0.15], p < 0.001), and Visual networks (β_std_ = 0.15, SE = 0.02, 95% CI [0.10, 0.19], p < 0.001; see Figure 2). In contrast, alignment negatively predicted exploration in the Limbic network (β_std_ = −0.09, SE = 0.02, 95% CI [−0.14, −0.05], p < 0.001; see Figure 2). For pairwise comparisons between slopes, see Supplementary Material 3. The covariate number of turns (β_std_ = 0.47, SE = 0.03, t = 17.32, p < .001) was also positively associated with neural exploration, whereas the trial number (β_std_ = 0.02, SE = 0.02, t = 0.84, p = .39) was not.

**Figure 2.**
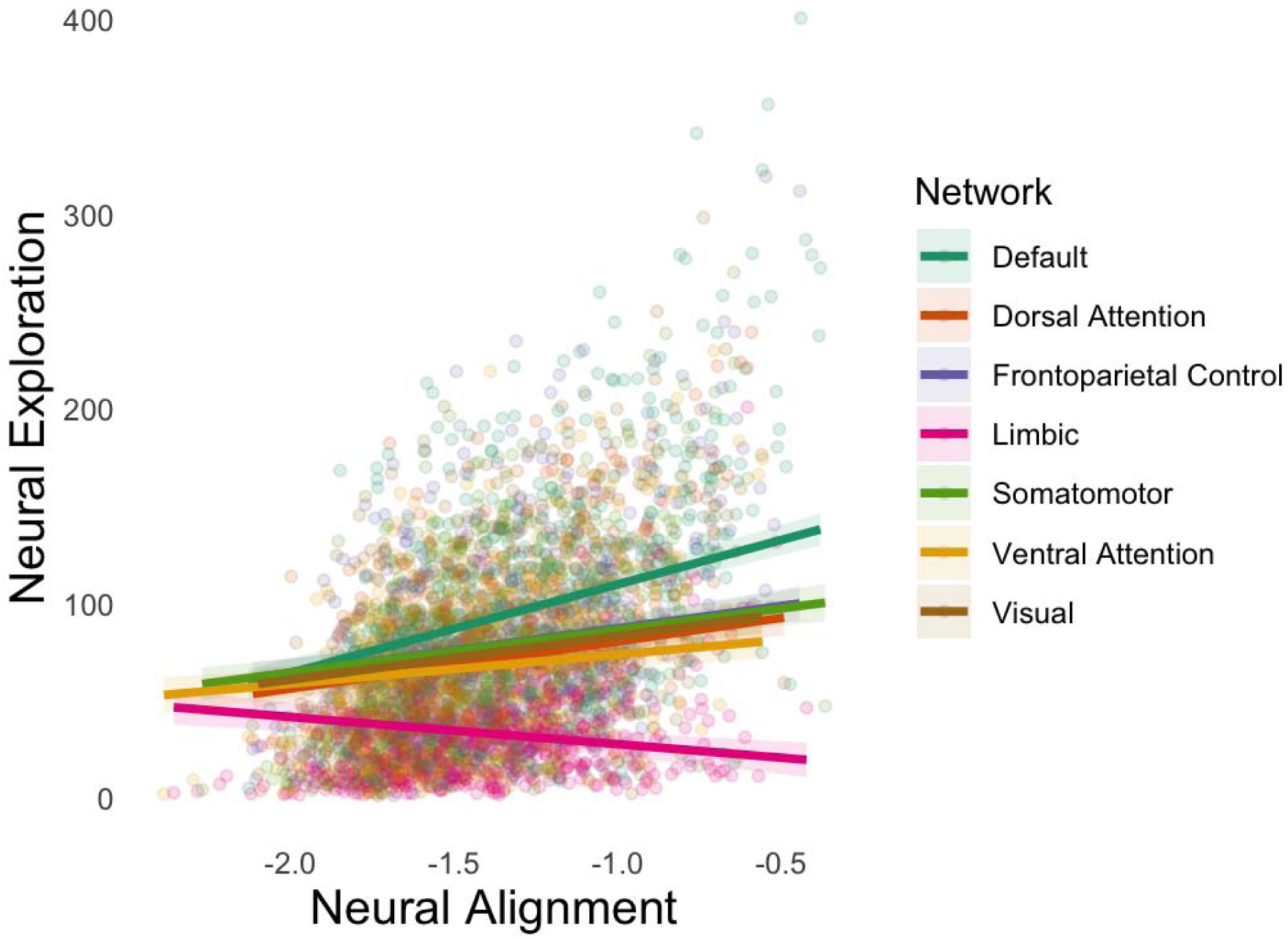
Dyads that are more aligned in representational space explore more in neural representational space across all networks except the limbic network. Fitted values and 95% confidence intervals were calculated using the model’s fixed effects and variance– covariance matrix.

### Dyads who exhibit more neural alignment and neural exploration also explore more conversational topics

This study derives novel neural measures of alignment and exploration from natural conversation using an HHMM. Because these measures capture abstract, latent neural dynamics, we sought to clarify how they relate to a more observable feature of conversation: the extent to which partners explore different topics. Prior studies have linked topic exploration to positive social outcomes, including greater social connection in the present dataset (Speer et al., 2024) and greater agreement in an independent dataset (Speer et al., 2026). Topic exploration may therefore provide a meaningful linguistic marker of exploratory conversational dynamics. Here, we test whether HHMM-derived neural alignment and exploration are associated with topic exploration.

Correspondence between these neural and linguistic measures would connect latent neural dynamics to overt conversational behavior and strengthen the interpretation and validity of the novel neural measures. We used topic modeling to quantify how much conversational content participants traversed (see Methods). Then, we used the same multilevel modeling approach as above to examine whether neural alignment and exploration are associated with topic exploration (See Methods). Model comparison revealed that the effect of representational alignment on topic exploration did not significantly differ across networks (χ²(6) = 6.68, *p* = .35), nor did the effect of neural exploration on topic exploration (χ²(6) = 3.19, p = .784). We therefore report results from the more parsimonious main-effects model.

Overall we see that both neural alignment and exploration predict topic exploration. Across networks, greater representational alignment positively predicted greater topic exploration (β_std_ = 0.041, SE = 0.015, t = 2.64, p = .008; see Figure 3), as did the number of conversational turns (β_std_ = 0.539, SE = 0.021, t = 26.306, p < .001) and the trial number (β_std_ = 0.11, SE = 0.013, t = 9.07, p < .001). Greater neural exploration *also* predicted greater topic exploration (β_std_ = 0.097, SE = 0.019, t = 5.02, p < .001; see Figure 3), as did the number of conversational turns (β_std_ = 0.518, SE = 0.021, t = 25.18, p < .001) and the trial number (β_std_ = 0.116, SE = 0.013, t = 9.88, p <.001). This suggests that both neural alignment and neural exploration are linked to observable conversation patterns, which demonstrates their external validity.

**Figure 3.**
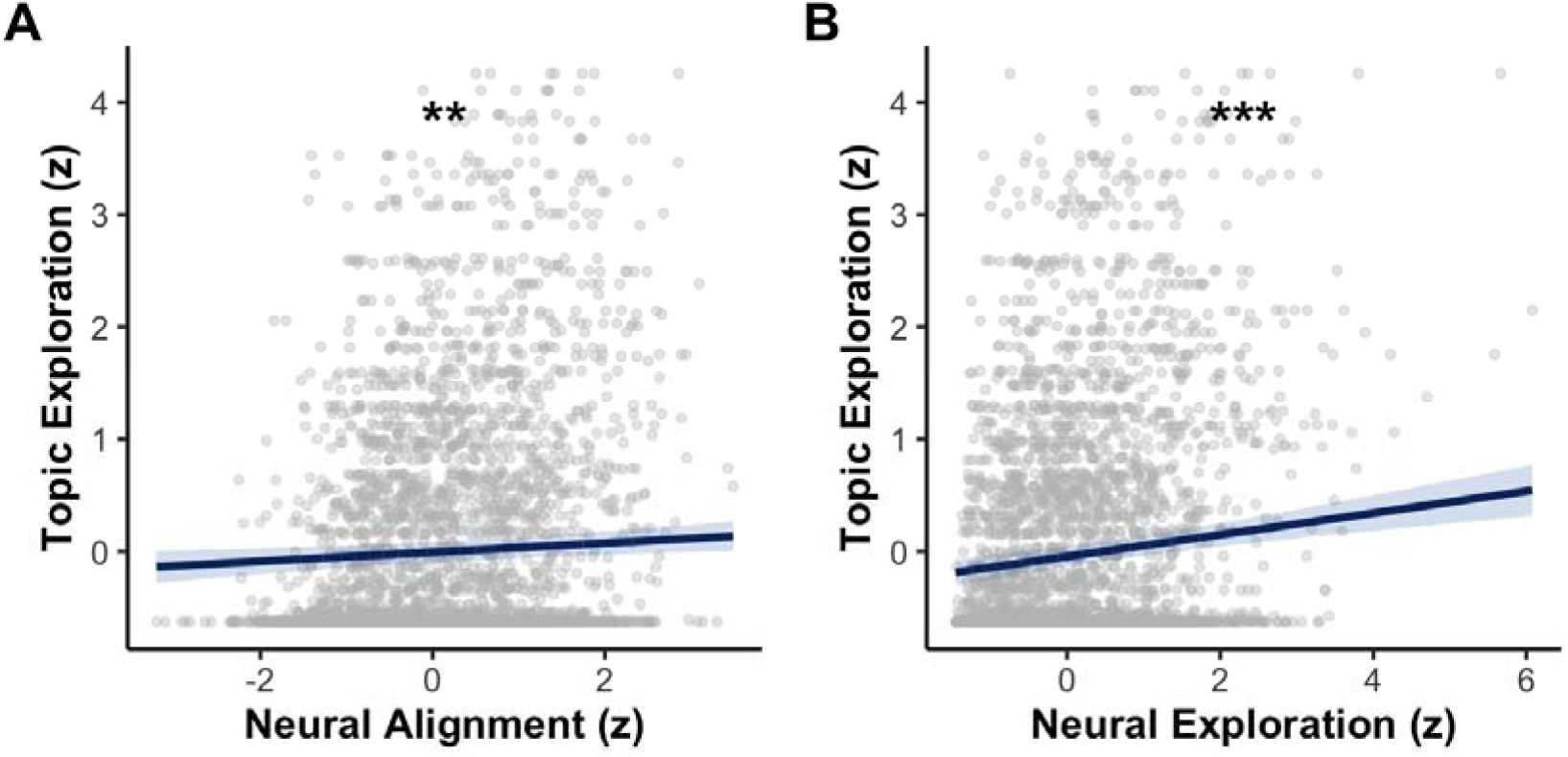
A) Dyads who are more aligned in neural representational space explore more topics across networks. B) Dyads who explore more neural representational space explore more topics. Fitted values and 95 % confidence intervals were calculated using the model’s fixed effects and variance–covariance matrix. † p < 0.10, * p < 0.05, ** p < 0.01, *** p < 0.005.

### Higher neural exploration, but not neural alignment, is associated with higher conversation quality

Because prior work has linked both alignment and exploration to successful conversations, we tested whether neural alignment and neural exploration each predicted self-reported conversation quality while controlling for social history (friends vs. strangers). As in previous studies (Mwilambwe-Tshilobo et al., 2025; Speer et al., 2024), conversation quality was captured along the dimensions of closeness, enjoyment, and similarity. We fit two separate models: one using alignment to predict conversation quality (closeness, enjoyment and similarity) and one using exploration. For the alignment and two of the exploration models, model comparisons indicated that associations with conversation quality did not differ across brain networks (all ps > 0.87) or as a function of social history (all ps > 0.14), so we report main-effects models (the same slope for all networks & both friend and stranger dyads). For the third exploration model, the association between exploration and similarity differed significantly between friends and strangers (F(1,387) = 4.38, p = 0.037). So we report an interaction model (separate slopes for friends and strangers) for similarity and main-effects models for closeness and enjoyment.

Overall, we see that exploration, but not alignment, predicts positive conversation quality.

### Alignment

Representational alignment did not predict any of the conversation outcomes: it did not significantly predict closeness (β_std_ = 0.035, SE = 0.058, t = 0.61, p = .543) or enjoyment (β_std_= 0.026, SE = 0.064, t = 0.40, p = .687), and was marginally associated with greater perceived similarity (β_std_ = 0.080, SE = 0.046, t = 1.74, p = .082; see Figure 4).

**Figure 4.**
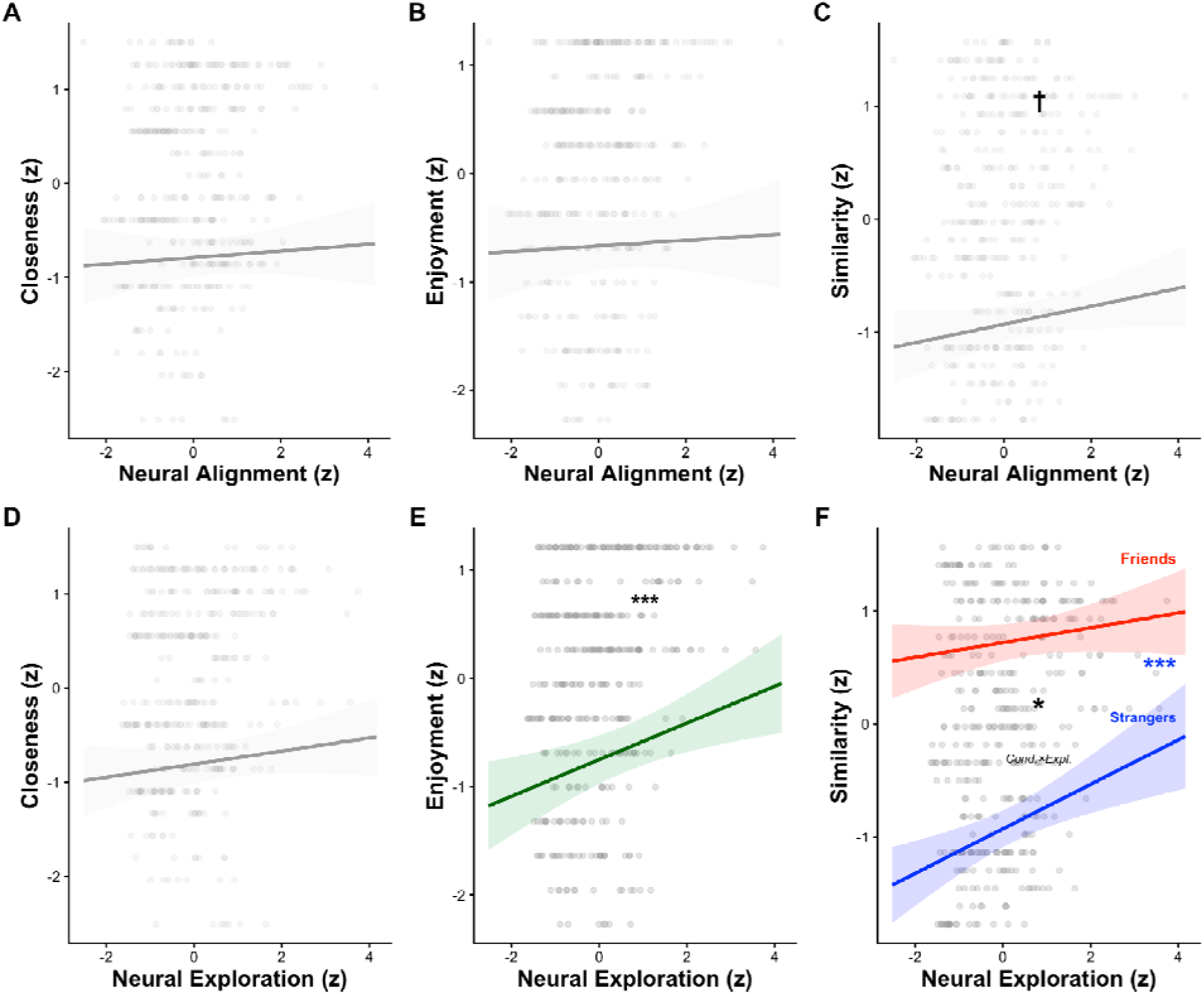
Dyads who explore more neural representational space have higher-quality conversations. Fitted values and 95% confidence intervals were calculated using the model’s fixed effects and variance–covariance matrix. Non-significant associations are shown with transparent grey regression lines and confidence ribbons; significant associations are shown in green. For similarity, a significant exploration × condition interaction was found: separate regression lines are shown for friends (red) and strangers (blue), with significance indicators reflecting simple slopes within each group. The central asterisk and subscript label (“Cond.×Expl.”) indicate the significance of the interaction term. † p < 0.10, * p < 0.05, ** p < 0.01, *** p < 0.005.

### Exploration

Neural exploration predicted two of the three conversation outcomes: it did not significantly predict closeness (β_std_ = 0.070, SE = 0.052, t = 1.35, p = .179), but it did predict greater enjoyment (β_std_ = 0.169, SE = 0.057, t = 2.96, p = .003, see Figure 4), and greater perceived similarity among strangers (simple slope: β*_std_* = 0.197, SE = 0.055, t = 3.56, p < .001), but not friends (simple slope: β*_std_* = 0.065, SE = 0.049, t = 1.34, p = .181). For perceived similarity, turns showed a marginal negative association in the alignment model (β*std* = −0.075, SE = 0.045, t = −1.66, p = .099) but not in the exploration interaction model (β*std* = −0.063, SE = 0.039, t = −1.62, p = .106).

### Joint Model

When neural alignment and neural exploration were entered into the same model to test their unique contributions to conversation quality, only neural exploration explained significant unique variance in enjoyment and similarity, whereas neither neural measure explained unique variance in closeness (see Supplementary Material 4).

### Friends represent conversational events with greater neural alignment than strangers

To test whether shared social history (i.e., friendship) was associated with greater neural alignment and exploration, as predicted by our shared-mental-map account, we compared friend and stranger dyads. To account for the non-independence of multiple trials per dyad, we fit mixed-effects models with condition (friends vs strangers) and network as fixed effects and a random intercept for dyad. We first compared a main-effects model to one that additionally included a Condition × network interaction. We report network-specific effects for both alignment and exploration because the interaction model marginally improved fit over the main-effects model for alignment (LRT χ*²*(6) = 10.54, *p* = .10), and significantly improved fit for exploration (LRT χ*²*(6) = 13.56, *p* = .035).

Overall, we see that friends show greater alignment and exploration than strangers.

### Alignment

Friends showed greater representational alignment than strangers in the frontoparietal control (β = 0.36, t(55) = 2.43, p = .019), dorsal attention (β = 0.31, t(55) = 2.11, p = .040), and limbic networks (β = 0.31, t(55) = 2.08, p = .042; see Figure 5). The default network showed a marginal difference (β = 0.27, t(55) = 1.83, p = .073; see Figure 5), whereas no other networks differed significantly between friends and strangers.

**Figure 5.**
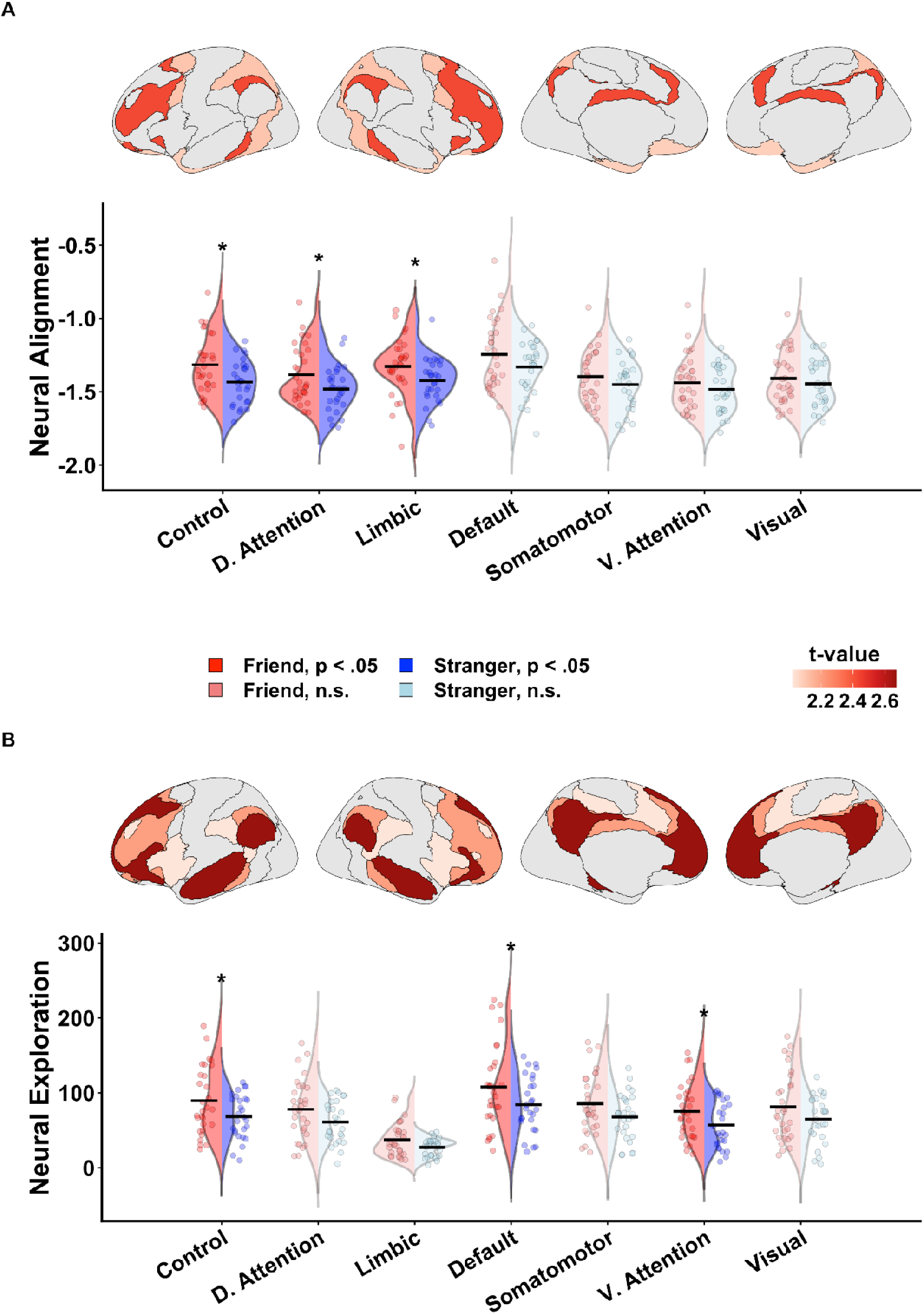
A) Friends represent conversational events with greater neural alignment than strangers. B) Friends explore more representational space than strangers. Top rows: t-values for networks with significant Friend vs. Stranger differences plotted on the brain surface, derived from per-network contrasts of a mixed-effects model including a Condition × network interaction (random intercept for dyad). Darker red colors indicate higher t-values. Bottom rows: Violin plots for all seven Yeo networks showing Friend (left, blue) and Stranger (right, red) distributions. Significant networks are shown in full color; non-significant networks are shown in muted colors. The dash indicates the group mean. * = p < .05.

### Exploration

Friends exhibited greater neural exploration than strangers in the default (β = 0.48, t(55) = 2.67, p = .010), frontoparietal control (β = 0.40, t(55) = 2.21, p = .032), and ventral attention networks (β = 0.36, t(55) = 2.03, p = .048; see Figure 5). Marginal differences emerged in the somatomotor (β = 0.36, t(55) = 1.99, p = .051; see Figure 5), dorsal attention (β = 0.34, t(55) = 1.91, p = .062, see Figure 5), and visual networks (β = 0.33, t(55) = 1.83, p = .073; see Figure 5). Together, these findings indicate that friends explored more representational space than strangers.

## Discussion

What happens in the brain during successful communication? Neuroscientists have proposed two neural mechanisms that may support good conversation: alignment and exploration. Although both have been linked to positive social outcomes (Doré & Morris, 2018; Nguyen et al., 2021; Parkinson et al., 2018b; Ravreby et al., 2022; Sened et al., 2024; Shen et al., 2025; Sievers et al., 2024; Speer et al., 2024, 2026; Street et al., 1983), they have mostly been studied separately, leaving open whether they compete, operate independently, or complement one another during real-time interaction. Using HHMM to quantify both dynamics in the same shared latent space, we found evidence for *coordinated exploration*: greater neural alignment was associated with greater neural exploration. This suggests that alignment may facilitate exploration, allowing people to move into new territory while remaining mutually oriented. Both measures were also linked to broader topic exploration. Notably, greater neural exploration but *not* alignment was associated with better conversation quality. Friends showed greater neural alignment and exploration than strangers. Together, these findings suggest that alignment and exploration are complementary rather than competing dynamics, and that exploration is a distinct neural dimension of conversation that is especially relevant to how the interaction is experienced.

Prior work has often framed alignment as a process of convergence, in which conversation partners become more similar in their behavior, language, physiology, or neural responses (Nguyen et al., 2021; Peng et al., 2026; Shaw et al., 2018; Sievers et al., 2024). Exploration, by contrast, has often been framed as movement into new conversational territory, implying novelty, divergence, and distance between partners (Burns et al., 2025; Speer et al., 2024, 2025, 2026). Our findings challenge this tension by showing that the two dynamics can coexist and complement each other in the form of *coordinated exploration* within the same conversation. We see that neural exploration and neural alignment are positively correlated in all networks but one. One possible explanation is that both dynamics are supported by an underlying shared mental map. Friends, for instance, share inside jokes, past experiences, common references, and topic knowledge. These overlapping semantic, autobiographical, affective, and social associations may help partners represent the same conversational events more similarly. At the same time, higher representational alignment of the current event prepares friends to make larger topic jumps to the next conversation event without losing each other. Thus, neural event alignment may be best understood as a readout of a shared mental map that allows participants to engage in *coordinated exploration* and cover a vast terrain of topics.

Because HHMM quantifies neural alignment and exploration in an *abstract* shared latent space, it is important to establish whether these measures correspond to observable features of conversation. We therefore used topic exploration as a linguistic and semantic index of conversational breadth. Dyads with greater neural alignment and exploration also covered a broader range of topics and these associations did not significantly differ across networks. This convergence between neural and linguistic measures provides evidence for the convergent validity of the HHMM-derived metrics. Neural alignment and exploration, therefore, capture behaviorally meaningful variation in conversational breadth, rather than merely reflecting geometric properties of the latent neural space.

After establishing that neural alignment and exploration complement each other and that both measures are related to observable conversation breadth, we next turned to conversation outcomes. The previous literature suggests that both alignment and exploration can support good conversations, albeit via different mechanisms. Alignment may support understanding and coordination (daSilva & Wood, 2024; Garrod & Pickering, 2004; Koban et al., 2019), whereas exploration may render the conversation more engaging, informative, and socially revealing (Burns et al., 2025; Speer et al., 2024, 2025, 2026; Templeton et al., 2024; Tsang et al., 2025). Previous analyses of the same conversations linked greater neural, linguistic, and topic exploration to better self-reported conversation quality (Speer et al., 2024). Here we had the opportunity to directly compare the contribution of neural alignment and neural exploration to these outcomes. Across three dimensions of self-reported conversation quality, namely closeness, enjoyment, and perceived similarity, neural exploration, but *not* neural alignment, predicted conversation success. Collectively, this suggests that even though alignment and exploration coexist and complement each other, exploration is not merely a downstream byproduct of alignment but a distinct neural dynamic of conversation that is especially important for predicting its social outcomes.

Coordinated exploration varied systematically with relationship context: Friends showed both greater event-level alignment and broader conversation-level exploration than strangers. This pattern is consistent with the hypothesis that shared social history increases the fidelity of the shared mental map that enables participants to represent conversational events more similarly and explore more broadly. Our findings also reframe results from a previous analysis of the same conversations, which showed that friends became more exploratory by diverging in neural, linguistic, and topic space, whereas strangers became more aligned by converging over time (Speer et al., 2024). The present HHMM analysis replicates friends’ greater exploration while revealing that this broad movement coexisted with greater similarity in partners’ representations. Thus, divergence between partners’ trajectories and the breadth of their exploration are not interchangeable: Partners can move broadly through representational space while remaining locally aligned. This distinction dovetails with recent fMRI hyperscanning work showing that novelty, operationalized as the amount of new mental-state or topic space a dyad covers, predicted agreement more strongly than divergence, or the degree to which partners moved away from each other (Speer et al., 2026).Together, these findings suggest that productive exploration may depend less on moving away from one’s partner than on jointly moving into new territory while maintaining a shared understanding of the current event.

The pattern of network-level results offers a tentative window into the systems that support coordinated exploration in conversation. Alignment predicted greater exploration across all networks except the limbic network. Friend–stranger differences were also concentrated in only partially overlapping systems: friends showed greater alignment in frontoparietal control, dorsal attention, and limbic networks, and greater exploration in default, frontoparietal control, and ventral attention networks. This pattern of results suggests that coordinated exploration may depend on multiple components: flexible control over conversational transitions (frontoparietal network; Cocuzza et al., 2020; Cole et al., 2013; Dosenbach et al., 2007), shared attention to the current event (dorsal attention network; Corbetta & Shulman, 2002; Vossel et al., 2014), reorienting toward new conversational content (ventral attention network; Corbetta et al., 2008), and movement through meaning-based representations such as memories, narratives, and perspectives (default network; Andrews Hanna et al., 2014; Baldassano et al., 2017; Schurz et al., 2021). The limbic network showed a different pattern. Friends were more aligned in this network, but greater limbic alignment predicted less exploration, suggesting that affective alignment may stabilize rather than expand affective state space. For example, when partners share a stable emotional tone or appraisal, they may remain affectively anchored even as the conversation moves through new semantic or attentional territory. Thus, coordinated exploration may involve broad representational movement supported by non-limbic systems, alongside a more stable shared affective frame. These interpretations remain tentative because assigning psychological functions to broad networks involves reverse inference (Poldrack, 2006).

Although the present findings suggest that coordinated exploration is associated with successful conversation, the current design and HHMM implementation impose three interpretative constraints. First, the analyses are correlational. Because we did not causally manipulate exploration or alignment, we cannot determine whether alignment promotes exploration, exploration promotes alignment, or they are both caused by a third variable such as shared history or common ground. Second, the HHMM implementation assumes that both conversation partners move through the same sequences of events, although they may enter and exit those events at different times. This assumption is useful for estimating shared event representations, but it may be less appropriate for less coherent or more asymmetric conversations, in which one partner skips events, lingers on prior events, or represents a different event altogether. Future work could relax this constraint by allowing partially shared, idiosyncratic, or partner-specific event sequences. Third, HHMM requires the average event length to be specified in advance, which determines the expected number of events per conversation. In the present study, this parameter was based on average conversational turn length. Although turns are a meaningful unit of dialogue, they may not always correspond to neural or conceptual events: a single turn may contain multiple events, and a single event may span several turns. Future work should allow the number and duration of events to be estimated more directly from the neural and linguistic timecourses. Collectively, these extensions would help investigate whether shared representational alignment causally supports coordinated exploration by using models that allow more flexible event sequences and would link HHMM-derived events more closely to the actual content and dynamics of conversation.

This study bridges alignment and exploration accounts of conversation by testing both dynamics within the same real-time interaction. Using HHMM, we separated local event-level alignment, how similarly partners represented the current conversational event, from broader conversation-level exploration, how widely dyads moved through shared neural space. This approach shows that alignment and exploration are not mutually exclusive: partners can remain coordinated while moving through a wider neural and conversational landscape. More broadly, the study demonstrates HHMM’s utility for modeling social interaction as a shared neural trajectory rather than as simple synchrony between partners. Good conversation may therefore be best understood as coordinated exploration: moving into new neural, conversational, and social territory while remaining grounded in mutual understanding and perspective taking.

## Methods

In this study, friend and stranger dyads engaged in naturalistic conversations while their brain activity was simultaneously recorded with fMRI scanners in adjacent rooms. We tested how alignment and exploration interact during conversation. In addition, we examined how these two processes relate to topic exploration and how they are associated with self-reported conversation quality. Lastly, we tested how exploration and alignment differ between friends and strangers.

## Participants

Sixty-three dyads (126 individuals) engaged in live conversations during fMRI hyperscanning. We removed seven pairs because one or both members had incomplete data due to technical issues in the conversational or preprocessing phases. Our final sample comprised 57 pairs (114 participants; M_age = 20.6 years, SD = 3.0; 73 women, 41 men, none identifying as non-binary). Thirty of those pairs were self-reported friends (they attested that they interact at least four days a week for at least 3 months), and 27 pairs of strangers were randomly matched and unacquainted before the study. All participants were at least 18 years old, and the protocol was approved by Princeton University’s Institutional Review Board.

## Conversation Task

Each dyad completed 10 conversation trials. On each trial, dyads engaged in a three-minute conversation prompted by questions from the “Fast Friends” procedure (Aron et al., 1997), which are designed to build rapport. Prompts increased in personal disclosure—from low (“Would you like to be famous? In what way?”) through medium (“What do you value most in a friendship?”) to high intimacy (“Share with your partner an embarrassing moment in your life.”). For each prompt, who spoke first was randomized, and speakers indicated the end of their turn by pressing a button to cue their partner.

Trials were organized into five runs; each run comprised two trials. A nine-second screen displayed the prompt, followed by 180 seconds of turn-based conversation (see Figure 6). Runs began and ended with a 12-second fixation cross. Each dyad completed 10 trials. Across the 570 dyad-trials, trials contained an average of 6.64 conversational turns (SD = 4.66, median = 5, IQR = 4-8, range = 2-41). Friend dyads produced more turns per trial on average (M = 7.73, SD = 5.77) than stranger dyads (M = 5.42, SD = 2.48). For details on speech recording, please refer to Supplementary Material.

**Figure 6.**
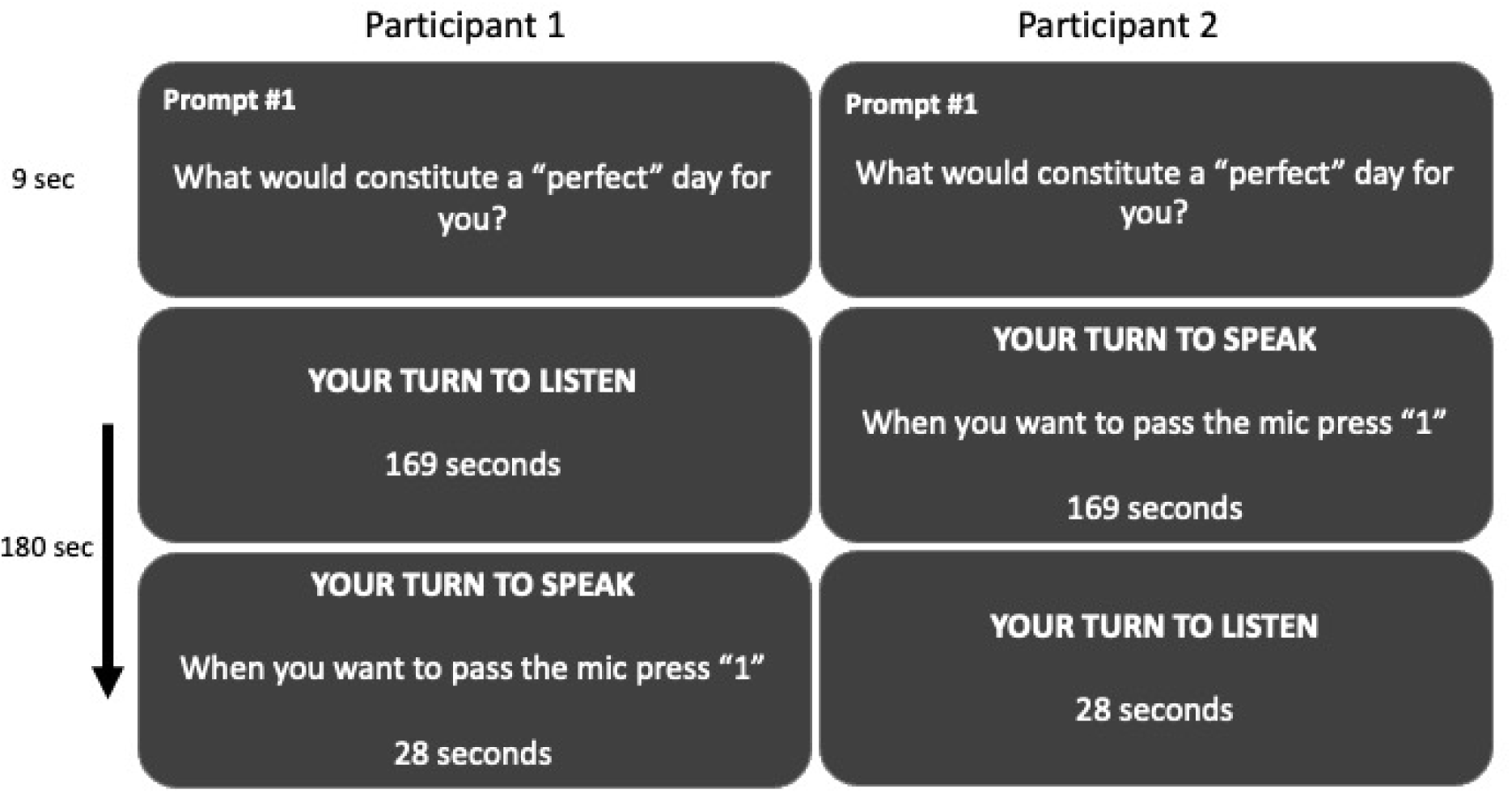
Illustration of a sample trial from each participant’s viewpoint. Both members of the dyad first see the discussion prompt for 9 seconds. They are randomly assigned the roles of speaker and listener, and the speaker begins talking. At any point, the speaker can press a button to transfer the speaking role to their partner. Throughout the trial, a timer displayed at the bottom of the screen indicates the remaining time.

## Imaging Procedure

### MRI Data Acquisition

Functional and structural scans were collected on 3T Siemens Skyra and Prisma systems using identical settings. Whole-brain functional images were acquired in an interleaved sequence with 3 mm slice thickness, 3 × 3 mm in-plane resolution, an 80° flip angle, TE = 28 ms, and TR = 1500 ms. For anatomical reference, a high-resolution T1-weighted scan was obtained (1 × 1 × 1 mm voxels, 176 sagittal slices, flip angle = 9°, TE = 2.98 ms, TR = 2300 ms). To limit head motion, participants’ heads were cushioned with foam padding.

### Imaging preprocessing

All fMRI scans were processed with fMRIPrep 20.2.0, a Nipype-based pipeline that automatically tailors its workflows to use state-of-the-art algorithms with minimal manual intervention (Speer et al., 2024, 2024; Zada et al., 2026). First, each T1-weighted volume was corrected for intensity nonuniformity and skull-stripped, then nonlinearly registered to the ICBM 152 Nonlinear Asymmetrical 2009c template and segmented into cerebrospinal fluid, white matter, and gray matter. Functional runs were distortion-corrected by coregistering them to their own T1w image, refining alignment with a nine-degree-of-freedom boundary-based registration, and then underwent motion correction, field distortion adjustment, and spatial normalization in one resampling step using Lanczos interpolation. Physiological noise components were extracted via CompCor (six temporal and six anatomical components). To further remove speaking-related motion artifacts and field distortions, we applied an optimized confound regression model that maximizes inter subject correlation detection to the preprocessed data. This model included head motion parameters (with their cosine transforms, first derivatives, and squared terms), white matter and CSF signals (plus derivatives), and task timing regressors. The resulting residual time series constituted our cleaned dataset, which we then segmented by prompt to examine trial-by-trial effects over time.

### Neural parcellation

We assessed network-specific model performance as in prior work on the same dataset (Mwilambwe-Tshilobo et al., 2025), by calculating the average model performance in the 7 resting-state functional networks (i.e., intrinsic networks) as defined by Yeo et al. (2011).

## Measures

### Conversation Quality

At the end of each conversation, participants completed a questionnaire assessing how socially connected they felt to their partner. To uncover underlying dimensions that summarize these measures, a previous study performed factor analyses (Speer et al., 2024). The items captured three latent dimensions of social connection: 1) how close the participant felt to their partner, 2) the enjoyment of the conversation, and 3) how similar the participant felt to their conversation partner (Speer et al., 2024). In the current study, we tested whether our neural measures (explained below) predict these conversation outcomes.

### Neural Measures

To test our hypothesis that shared spatial representations of conversational events enable friends to engage in more exploratory exchanges, we employed Hyper Hidden Markov Modeling (HHMM) to project each partner’s neural states into a common latent space and segment them into meaningful events. This approach allows us to characterize how each dyad represented the conversation in a shared latent space. In this study, we use the latent-space representations of each event to track the neural dynamics in each participant.

#### Hyper-HMM

We employed Hyper HMM (Lee et al., 2023), which jointly aligned temporal and spatial features across a dyad into a shared low-dimensional event space (see Figure 7). The model made three assumptions: (i) The conversation proceeds through a series of *events* (e.g., topic shifts, questions/answers, turns), averaging 6.64 turns per 180-s conversation, or approximately 18 TRs per turn. Both partners pass through the same sequence. However, the time spent in each event is allowed to differ across participants. Stated differently, while both partners proceed through the same events their event boundaries are allowed to occur at different timepoints (see Figure 7). (ii) Each event is represented by a vector in the shared latent space (see diamonds representing events in Figure 2). Each vector reflects a neural representation of what was being discussed at that moment (e.g., recognizing that the speaker just asked a question, or that the dyad shifted topics). (iii) Partners are linked by their individual linear transformation matrices, which map spatial patterns of activity in regions to a shared latent space. This transformation accounts for the fact that the same neural state during a conversation may appear as a different voxel pattern in each partner. The geometry of this space encodes the similarity between events (for similarity between events see green arrows in Figure 2). Events that are close together are more similar than events that are more distant. When two partners align on an event (e.g., “discussing the weather” or “asking a question”), their corresponding vectors will be close in latent space, reflecting joint attention, shared affective states, and predictive processing in conversation. We determined the dimensionality of this space via a grid search over 2 to 10 dimensions and ridge penalties based on the length of the scans. Two dimensions and a ridge penalty of α = 1000 minimized log-likelihood on pilot neural data (for details see Supplementary Material 2).

**Figure 7.**
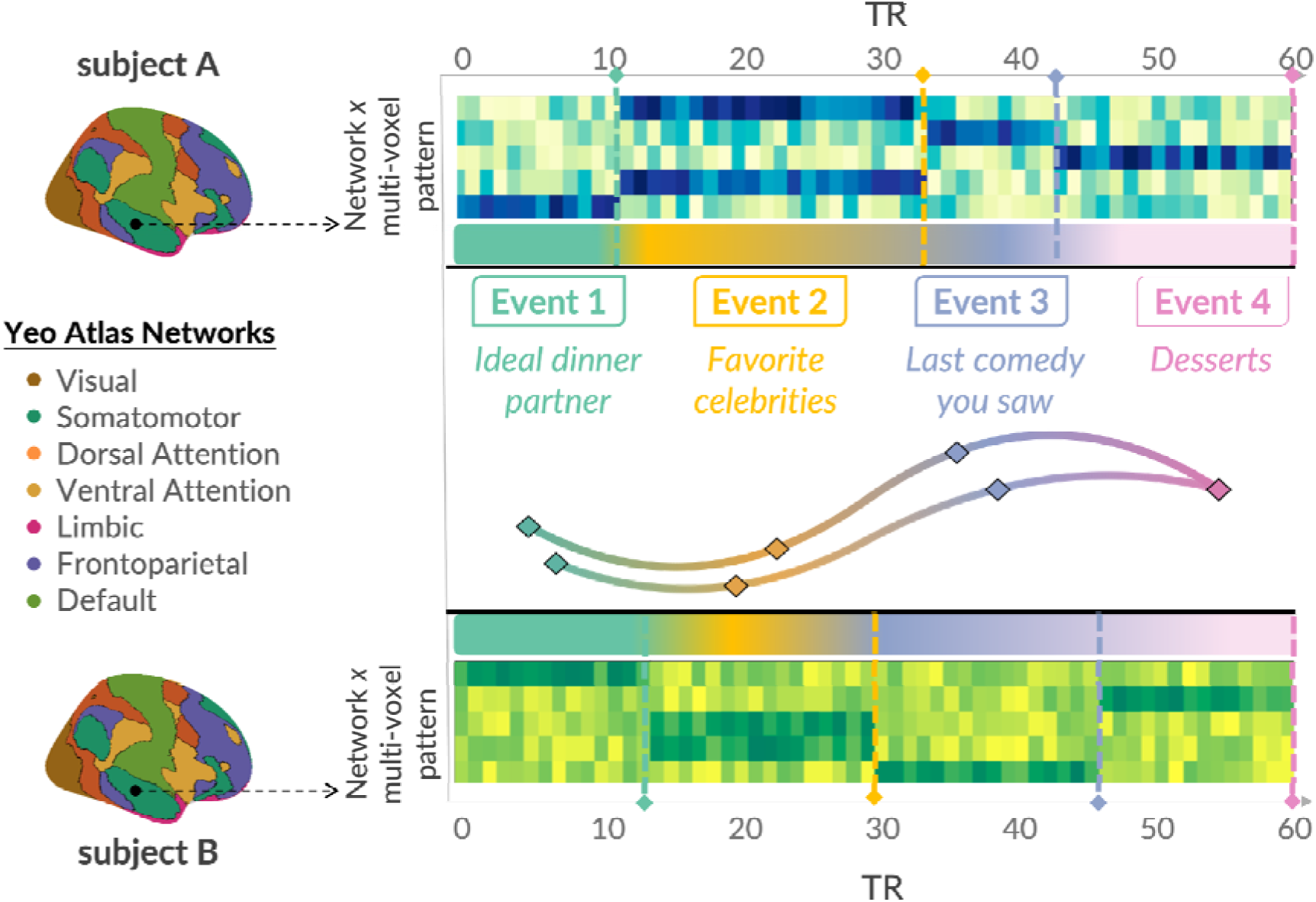
*Measuring learned event representations from the conversation partners’ neural data.* The model is fit to voxel patterns from both conversation partners (partner 1’s pattern in blue and partner 2’s pattern in green), successfully learning a temporal alignment into corresponding events despite temporal and representational differences between the two conversation partners.

We applied the same dimensionality to all dyads because the goal was to estimate a common representational space in which participants can be compared directly (Guntupalli et al., 2016; Haxby et al., 2011, 2020). Using the same number of dimensions across participants is necessary for distances, trajectories, and event patterns to be interpreted in a common coordinate system; allowing each dyad to have a different number of dimensions would undermine direct comparison. This same convention is also used when hyperalignment is applied to other naturalistic datasets, where movie-or story-evoked responses are used to learn a single reduced-dimensional shared space plus participant-specific mappings into that space (Busch et al., 2021; Nastase et al., 2020).

We applied Hyper HMM to our fMRI conversation data. This fitting process alternated between inferring the timing of each partner’s neural events and refining both the shared latent space and the individual projection matrices. At each iteration, the neural pattern in each parcel was projected into the current two-dimensional latent space. A forward–backward algorithm (Baldassano et al., 2017) estimated the probability of each event over time. These estimates were used to refine partner-specific neural representations of each event. Next, we updated the shared latent space by combining all event representations across partners and extracting the top two principal components. Partner-specific projection matrices were re-estimated via ridge regression to map neural activity into this updated space. This process was repeated until model fit converged as evaluated by the log-likelihood. The result was a unified segmentation of the conversation into neural events that aligned across partners and a common latent space where each partner’s neural patterns co located via the learned transformation matrices.

For each dyad, we employed Hyper-HMM to segment each partner’s preprocessed neural time series into latent conversation events (see Figure 7). Hyper-HHM requires that we set a parameter setting the average event length. We chose the average length of a turn, calculated separately for each trial, because this was the smallest meaningful temporal unit of analysis in our conversation dataset at which content and roles can change. This choice implicitly assumes that each turn constitutes a new conversational event. This assumption is consistent with our topic-modeling approach, in which each turn is treated as a separate document that can be assigned a topic. Accordingly, a new conversational event may reflect either the introduction of a new similar or dissimilar topic or the continuation of the same topic by the other conversation partner. We conducted these analyses separately for each of the ten trials, for each of the 7 neural networks.

#### Neural alignment

We quantified how close partners’ representations of the same conversation event were in latent space. Specifically, for each event index identified by the HHMM, we measured the Mahalanobis distance between partner 1’s and partner 2’s neural state corresponding to that event (e.g., event 1 with event 1, event 2 with event 2). The Mahalanobis distance was chosen due to its insensitivity to the scale of the variables entered and because it removes redundant information from correlated variables. This distance between concurrent events captures how similarly the two partners represent the ongoing conversation, as indicated by the proximity of their neural patterns in representational space. We termed this measure *neural alignment*. We then averaged these event-wise distances across the conversation to obtain a dyad-level estimate of alignment. Using this measure, we tested how friends differ from strangers in how they represented conversational events in shared latent space.

#### Neural exploration

To measure neural exploration, we used the convex hull approach (Hart et al., 2018; Staerman et al., 2020). Specifically, we computed the area over the set of dyad-level event representations in the two-dimensional HHMM latent space. Each event contributed one joint dyadic point to the point cloud, such that □_□_ {□_□.□_, □ □^2^□=1,…, □_□_}where *z*_d,e_ denotes the two-dimensional latent-space representation of event *e* for dyad *d*, and *E_d_* is the number of dyad-level events. We then identified the smallest convex polygon containing all points in *P*_d_. Conceptually, this approach is like stretching a tight rubber band around all the pins on a board. Listing the ordered vertices of this convex polygon as □_□, 1,_ □_,□,□□_ where *h_d,k_* = (□_□,□_,_1_, □,_□ □,2_) the area was computed using the shoelace formula:

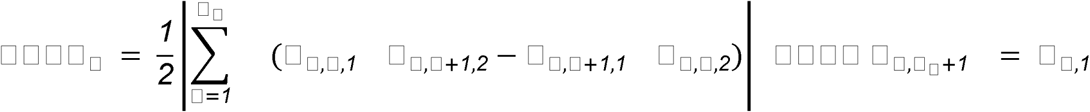

Where *h_d,k_* refers to the Cartesian coordinates of the ordered convex-hull vertices in two-dimensional latent space. This allows us to quantify the spread and distribution of conversation events. A high spread of conversation events may represent more flexible, wide-ranging exploratory conversations.

#### Topic Exploration

We also measured the amount of content space dyads covered in their conversation using topic modeling. Topic embeddings were computed in a previous study (Speer et al., 2024) using BERTopic, such that each turn was assigned a topic embedding. We projected the high-dimensional (N = 384) topic embeddings into two dimensions using t-distributed Stochastic Neighbor Embedding (t-SNE;van der Maarten & Hinton, 2008). All subsequent analyses were performed in this two-dimensional t-SNE space. To quantify how dispersed each trial’s topic points were in t-SNE space, we grouped observations by dyad and trial, extracted the t-SNE coordinates, and, when at least three points were present, computed the two-dimensional convex hull area (as explained above); trials with fewer than three points were assigned an area of zero.

## Statistical analyses

### General modeling approach

We used linear mixed-effects models to test condition effects (friends vs strangers), associations among neural exploration, neural alignment and topic exploration, and links between the neural measures and conversation quality. Continuous predictors, outcomes, and covariates were z-scored across the full dataset prior to analysis to facilitate interpretation of effect sizes. Models were fit using maximum likelihood (ML) for model comparison and restricted maximum likelihood (REML) for final parameter estimation. For each analysis, we first fit a main-effects model and then tested whether adding the theoretically relevant interaction improved model fit using nested model comparisons. In the main-effects model, the association between the predictor and outcome was constrained to be the same across levels of the moderator; in the interaction model, this association was allowed to vary, yielding separate slopes for each network or condition. Interaction terms were retained only when they significantly improved fit; otherwise, the more parsimonious main-effects model was retained. Models included random intercepts for dyads, with trials nested within dyads when supported by the data structure. When the richer random-effects structure led to non-convergence or singular fit, we used the simpler dyad-level random-intercept structure.

### Testing the association between neural alignment, neural exploration and topic exploration

We used the modeling framework described above to examine associations among neural alignment, neural exploration, and topic exploration at the trial level. For each model, we included the neural or topic-level predictor of interest, network, number of conversational turns, and trial number as fixed effects.

We fit three model sets: first, models predicting neural exploration from neural alignment; second, models predicting topic exploration from neural alignment; and third, models predicting topic exploration from neural exploration.

### Testing the effect of neural alignment and neural exploration on conversation quality

Because conversation-quality outcomes were measured at the dyad level, neural alignment and neural exploration were aggregated to the dyad level before analysis. For each neural measure, we first averaged values across events within each dyad × trial × network to obtain trial-level estimates, and then averaged those trial-level estimates across trials to create a dyad × network dataset. This two-step aggregation ensured that each trial contributed equally to the dyad-level score rather than allowing trials with more events to exert disproportionate influence.

Dyad-level outcomes included closeness, enjoyment, and perceived similarity from prior analyses of the same dataset (see Measures: Conversation Quality). For each outcome, we fit models with either neural alignment or neural exploration as the primary predictor separately, controlling for mean number of events, network, and condition. Condition was included because prior work on this dataset showed that friends reported greater enjoyment, similarity, and closeness (Speer et al., 2024). We also entered both alignment and exploration into the same model to see whether both explain unique parts of the variance in conversation quality.

### Testing the effect of friendship on neural alignment and exploration

To test the effect of social history on neural alignment and exploration, we fit separate mixed-effects models predicting neural alignment and exploration from condition, network, and their interaction using the modeling framework described above.

## Acknowledgments

This work was supported by NIMH grant R21MH127284 and PNI Research Innovator Award to Diana I. Tamir. This publication was supported by the Princeton University Library Open Fund Access Fund. The authors thank Zaid Zada for their feedback on earlier versions of this manuscript, and members of the Princeton Social Neuroscience Laboratory for helpful discussions. Funding for the acquisition of the data was provided in part by the Regina and John Scully ’66 Center for the Neuroscience of Mind and Behavior.

