## Supplementary material for "fMRI hyperscanning reveals that coordinated exploration predicts successful conversation"

**Supplementary Material 1 - Speech Recording**

All conversations during the fMRI sessions were captured with a dedicated MR‑safe recorder (FOMRI II; Optoacoustics Ltd). It uses two optical microphones set perpendicular to each other—one devoted solely to ambient sounds (the reference), the other picking up both ambient noise and the participant’s speech (the signal). A dual‑adaptive filter then employs a least‑mean‑squares algorithm to subtract the reference from the signal. To keep that subtraction on track, the reference channel itself is adaptively filtered, with its gain tweaked continuously based on the leftover noise and the original reference input. A built‑in voice activity detector is incorporated during speech to prevent the filter from diverging, and an additional spectral enhancement filter polishes the output in real time. Participants spoke and listened via an audio link bridged between the two adjacent rooms.

**Supplementary Material 2 - Determining Hyperparameters for HHMM**

To identify optimal hyperparameters for the dyadic HHMM, we implemented a grid search across three pilot dyads that were not included in the main analysis and across a set of predefined regions of interest (ROIs). The ROIs examined were anterior medial prefrontal cortex (anterior MPFC), temporoparietal junction–angular gyrus (TPJ angular gyrus), posterior cingulate cortex/precuneus (PCC precuneus), temporoparietal junction–supramarginal gyrus (TPJ SMG), ventromedial prefrontal cortex (vmPFC), nucleus accumbens (NAcc), primary auditory cortex, primary visual cortex (V1), and superior temporal sulcus (STS).

For each dyad, preprocessed ROI time series from both partners were concatenated across conversational runs and restricted to task-relevant periods. A HHMM was then fit jointly to both participants’ neural data, assuming a common latent event sequence expressed in participant-specific voxel patterns.

We systematically varied two key hyperparameters: the dimensionality of the shared latent space (2, 3, 4, 5, 6, 8, 10) and the ridge regularization strength applied to voxel weights (1, 10, 100, 1000). Across pilot dyads and ROIs, the best-performing hyperparameter combination was a shared dimensionality of 2 and a ridge regularization parameter (α) of 1000. This consensus parameter set yielded an average log-likelihood of −162.31 and was therefore carried forward to the main analyses.

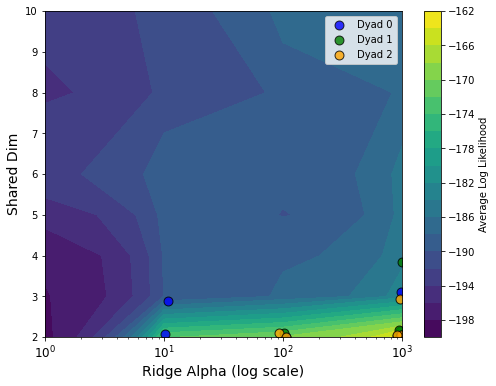

Figure S1. Grid search results for dyadic HHMM hyperparameters. The contour plot shows average log-likelihood across pilot dyads and selected ROIs as a function of shared latent dimensionality and ridge regularization strength (log-scaled). Colored points indicate the best-fitting parameter combination for each dyad–ROI pair. Higher values (brighter colors) reflect better model fit.

**Supplementary Material 3 - Pairwise Comparisons between the slopes for the model predicting neural exploration from neural alignment**

Pairwise comparisons of slopes, corrected for multiple comparisons using the Benjamini–Hochberg false discovery rate procedure, indicated that the alignment–exploration relationship was strongest in the Default network, where it was significantly stronger than in all other networks: Dorsal Attention (p < .001), Frontoparietal Control (p < .001), Somatomotor (p < .001), Ventral Attention (p < .001), Visual (p < .001), and Limbic (p < .001). In contrast, the relationship was weakest in the Limbic network, where it was significantly weaker than in the Dorsal Attention, Frontoparietal Control, Somatomotor, Ventral Attention, and Visual networks (all ps < .001). No other pairwise differences were significant after correction.

**Supplementary Material 4: Joint Models Including Neural Alignment and Neural Exploration**

Because our analysis shows that neural alignment is associated with neural exploration, we conducted supplementary analyses in which both predictors were entered into the same model. These models tested whether each neural measure explained unique variance in self-reported conversation quality while controlling for the other neural measure, condition, brain network, and number of turns.

For each outcome, closeness, enjoyment and similarity, we fit a joint model including neural alignment, neural exploration, number of conversational events, brain network, and condition. We then compared this model to reduced models that omitted either alignment or exploration.

For closeness, neither neural alignment nor neural exploration explained unique variance. Adding alignment to a model that already included exploration did not improve model fit (F(1, 387) = 0.08, p = 0.784). Adding exploration to a model that already included alignment also did not significantly improve fit (F(1, 387) = 1.51, p = 0.220). In the joint model, neither alignment, (β_std_ = 0.016, SE = 0.059, t = 0.27, p = 0.784), nor exploration (β_std_ = 0.066, SE = 0.054, t = 1.23, p = 0.220), significantly predicted closeness.

For enjoyment, neural exploration remained a significant unique predictor, whereas neural alignment did not. Adding alignment to a model that already included exploration did not improve model fit (F(1, 387) = 0.13, p = 0.718). By contrast, adding exploration to a model that already included alignment significantly improved model fit (F(1, 387) = 8.71, p = 0.003). In the joint model, exploration positively predicted enjoyment (β_std_ = 0.175, SE = 0.059, t = 2.95, p = 0.003), whereas alignment did not (β_std_ = −0.024, SE = 0.065, t = −0.36, p = 0.718).

For perceived similarity, neural exploration again explained unique variance, whereas neural alignment did not. Adding alignment to a model that already included exploration did not improve model fit (F(1, 387) = 1.10, p = 0.296). Adding exploration to a model that already included alignment significantly improved model fit (F(1, 387) = 6.42, p = 0.012). In the joint main-effects model, exploration positively predicted perceived similarity, β_std_ = 0.108, SE = 0.043, t = 2.53, p = 0.012), whereas alignment did not (β_std_ = 0.049, SE = 0.047, t = 1.05, p = 0.296).

We also tested whether these joint-model associations differed by brain network or condition. Neither alignment nor exploration interacted with brain network for any outcome, all ps > .74. Alignment did not interact with condition for any outcome, all ps > .12. However, for perceived similarity, the association between exploration and similarity differed by condition, F(1, 386) = 3.89, p = .049. Simple slopes indicated that greater neural exploration predicted greater perceived similarity among strangers, βstd = 0.185, SE = 0.057, t = 3.21, p = .001, but not among friends, βstd = 0.059, SE = 0.049, t = 1.20, p = .230.

Together, these supplementary joint models suggest that the outcome associations reported in the main text are primarily attributable to neural exploration rather than neural alignment. Neural exploration uniquely predicted enjoyment and perceived similarity after accounting for alignment, whereas alignment did not uniquely predict any conversation-quality outcome after accounting for exploration.

**Supplementary Material 5 - Robustness check: Controlling for the effect of modelfit**

To test the robustness of our findings, we conducted additional analysis to test to what extent model fit impacts our results.

*Differences in model fit between friends and strangers.* As a first step, we tested whether there were significant differences in model fit between friends and strangers. To this end, we tested whether trial-level log-likelihood (HMM model fit quality) differed between conditions across the seven networks. A likelihood-ratio test comparing a main-effects model to a model including a Condition × network interaction indicated that the interaction significantly improved fit (LRT: Δdf = 6, ΔlogLik = 15.17, *p* < .0001), and the interaction model was retained for inference. Despite the significant interaction, no individual network showed a statistically significant Friend–Stranger difference at the per-network level. This indicates that trial-level log-likelihood does not differ systematically between Friend and Stranger dyads, and that subsequent analyses controlling for trial-level log-likelihood are not confounded by differential HMM fit across conditions.

*Testing the effect of model fit on neural alignment*. Next, we tested whether trial log-likelihood predicted neural alignment within each network, using a mixed-effects model with nested random effects (trials within dyads). As before, a likelihood-ratio test comparing a main-effects model to one including a trial log-likelihood × network interaction indicated that the interaction significantly improved fit (LRT: Δdf = 6, ΔlogLik = 7.26, *p* = .024), and the interaction model was retained. Per-network slopes estimated from the interaction model indicated a significant positive relationship between trial log-likelihood and neural alignment in four of seven networks: Default (*b* = 0.164, SE = 0.034, *t*(3407) = 4.82, *p* < .001), Dorsal Attention (*b* = 0.118, SE = 0.032, *t*(3407) = 3.64, *p* < .001), Limbic (*b* = 0.120, SE = 0.055, *t*(3407) = 2.20, *p* = .028), and Somatomotor (*b* = 0.075, SE = 0.035, *t*(3407) = 2.17, *p* = .030). The relationship did not reach significance in the Frontoparietal Control (*b* = 0.044, *p* = .191), Ventral Attention (*b* = 0.044, *p* = .169), or Visual (*b* = 0.014, *p* = .711) networks. These results indicate that trials with better model fit (higher log-likelihood) tend to exhibit greater between-brain alignment.

*Testing the effect of model fit on neural exploration*. Next, we tested whether trial log-likelihood predicted neural exploration within each network using the same nested mixed-effects structure as Analysis 1. The trial log-likelihood × network interaction model substantially outperformed the main-effects model (LRT: Δdf = 6, ΔlogLik = 111.50, *p* < .0001), and was retained for inference. In contrast to the alignment analysis, trial log-likelihood positively predicted neural exploration in all seven networks (all *p* ≤ .022). Slopes were large and consistent across most networks: Default (*b* = 0.571, SE = 0.019, *t*(3407) = 30.73, *p* < .001), Frontoparietal Control (*b* = 0.430, SE = 0.018, *t*(3407) = 23.42, *p* < .001), Visual (*b* = 0.413, SE = 0.020, *t*(3407) = 20.69, *p* < .001), Somatomotor (*b* = 0.398, SE = 0.019, *t*(3407) = 20.98, *p* < .001), Dorsal Attention (*b* = 0.373, SE = 0.018, *t*(3407) = 21.00, *p* < .001), and Ventral Attention (*b* = 0.363, SE = 0.018, *t*(3407) = 20.64, *p* < .001). The Limbic network showed a significant but substantially weaker relationship (*b* = 0.069, SE = 0.030, *t*(3407) = 2.30, *p* = .022), which drove the significant interaction term (trial log-likelihood × Limbic: *b* = −0.502, *t*(3407) = −14.61, *p* < .001). Overall, these results indicate that trials with higher-quality HMM state structure are associated with greater neural exploration.

*Group differences in neural alignment between friends and strangers, controlling for model fit.*

We tested whether neural alignment differed between Friend and Stranger dyads across networks, controlling for trial log-likelihood. A likelihood-ratio test indicated that adding a Condition × network interaction did not significantly improve model fit (LRT: Δdf = 6, ΔlogLik = 4.57, *p* = .166), and the main-effects model was retained for inference. In the main-effects model (REML), Friend dyads showed marginally higher neural alignment than Stranger dyads averaged across all networks (*b* = 0.227, SE = 0.126, *t*(55) = 1.79, *p* = .078). This effect did not reach conventional significance but is consistent in direction with greater neural coupling in Friend dyads. To more closely compare these results with the original analysis, we also examined per-network Friend vs. Stranger contrasts from the interaction model. Per-network estimates were broadly consistent with the original findings: the largest and most reliable effects emerged in the Frontoparietal Control (*b* = −0.333, SE = 0.143, *t*(55) = −2.336, *p* = .023) and Limbic (*b* = −0.330, SE = 0.143, *t*(55) = −2.317, *p* = .024) networks, both showing significantly greater alignment in Friend than Stranger dyads (negative estimates indicate Stranger < Friend). The Dorsal Attention network showed a marginal effect in the same direction (*b* = −0.265, SE = 0.143, *t*(55) = −1.858, *p* = .068), while the Default (*p* = .110), Somatomotor (*p* = .247), Ventral Attention (*p* = .303), and Visual (*p* = .436) networks did not reach significance. This per-network pattern mirrors the pattern observed in the analysis in the main text, suggesting the group difference in neural alignment is consistent across models.

*Group differences in neural alignment between friends and strangers, controlling for model fit.* We also tested whether neural exploration differed between Friend and Stranger dyads across networks, controlling for trial log-likelihood. A likelihood-ratio test indicated that the Condition × network interaction did not significantly improve model fit over the main-effects model (LRT: Δdf = 6, ΔlogLik = 4.82, *p* = .141), and the main-effects model was retained for inference. In the main-effects model, Friend dyads showed significantly greater neural exploration than Stranger dyads averaged across all networks (*b* = 0.295, SE = 0.132, *t*(55) = 2.241, *p* = .029). Trial log-likelihood was a strong positive predictor of neural exploration (*b* = 0.445, SE = 0.012, *t*(3926) = 36.826, *p* < .001). To more closely compare these results with the original analysis, we also examined per-network Friend vs. Stranger contrasts from the interaction model. The pattern was broadly consistent with the original findings and showed a relatively uniform Friend > Stranger effect across networks. Significant effects emerged in the Default (*b* = −0.347, SE = 0.139, *t*(55) = −2.492, *p* = .016), Frontoparietal Control (*b* = −0.343, SE = 0.139, *t*(55) = −2.466, *p* = .017), Somatomotor (*b* = −0.358, SE = 0.139, *t*(55) = −2.576, *p* = .013), and Ventral Attention (*b* = −0.313, SE = 0.139, *t*(55) = −2.248, *p* = .029) networks. The Visual (*p* = .059) and Limbic (*p* = .094) networks showed marginal effects in the same direction, and the Dorsal Attention network did not reach significance (*p* = .154). The absence of a significant Condition × network interaction, combined with the broad distribution of effects across networks, indicates that Friend dyads show greater neural exploration than Stranger dyads in a largely network-general manner.

*Neural alignment predicts neural exploration, controlling for model fit.* We tested whether trial-level neural alignment predicted neural exploration across networks, controlling for number of events, trial number, and trial log-likelihood. A likelihood-ratio test revealed that the alignment × network interaction significantly improved model fit over the main-effects model (LRT: Δdf = 6, ΔlogLik = 104.35, *p* < .001), and the interaction model was retained for inference. Per-network slopes from the interaction model indicated that the positive relationship between alignment and exploration was significant across all networks, with one exception. The Default network showed the strongest positive slope (*b* = 0.255, SE = 0.016, *t*(3406) = 15.643, *p* < .001), and significant positive slopes were also observed in the Frontoparietal Control (*b* = 0.141, SE = 0.018, *t*(3406) = 7.621, *p* < .001), Somatomotor (*b* = 0.097, SE = 0.019, *t*(3406) =

5.118, *p* < .001), Visual (*b* = 0.058, SE = 0.020, *t*(3406) = 2.954, *p* = .003), Dorsal Attention (*b* = 0.056, SE = 0.020, *t*(3406) = 2.840, *p* = .005), and Ventral Attention (*b* = 0.045, SE = 0.020, *t*(3406) = 2.281, *p* = .023) networks. In contrast, the Limbic network showed a significant *negative* slope (*b* = −0.061, SE = 0.018, *t*(3406) = −3.443, *p* = .001), indicating that greater alignment in Limbic regions was associated with *less* neural exploration. This closely mirrors the results in the main text, demonstrating the robustness of our effects.

*Neural alignment predicts topic exploration, controlling for model fit.* We tested whether trial-level neural alignment predicted topic exploration, controlling for number of turns, trial number, and trial log-likelihood. Because topic exploration is a trial-level measure (constant across networks within a trial), random effects were specified as a random intercept for dyad only. A likelihood-ratio test indicated that adding an alignment × network interaction did not significantly improve model fit (LRT: Δdf = 6, ΔlogLik = 3.27, p = .366), and the main-effects model was retained for inference. In the main-effects model, greater neural alignment significantly predicted greater topic exploration (b = 0.041, SE = 0.016, t(3756) = 2.519, p = .012). These results closely match the results in the main text, pointing to the robustness of our findings across different models.

*Neural exploration predicts topic exploration, controlling for model fit.* We tested whether trial-level neural exploration predicted topic exploration, combining observations across networks and controlling for number of conversation turns, trial number, and trial log-likelihood. As in the analysis above, random effects were specified as a random intercept for dyad only, given that topic exploration is invariant across networks within a trial. A likelihood-ratio test indicated that the neural exploration × network interaction did not improve model fit (LRT: Δdf = 6, ΔlogLik = 1.66, *p* = .767), and the main-effects model was retained. In the main-effects model, greater neural exploration significantly predicted greater topic exploration (*b* = 0.135,

SE = 0.025, *t*(3756) = 5.336, *p* < .001). This again dovetails with the effects reported in the main text, highlighting the robustness of our findings across models.

*Neural alignment does not predict conversation outcomes, controlling for model fit.* We tested whether mean trial-level neural alignment predicted post-scan ratings of closeness, enjoyment, and perceived similarity, using data aggregated to the dyad × network level. For each outcome, we fit a linear regression with mean alignment, number of events, trial log-likelihood, network, and dyad type (Condition) as predictors, and tested two additional interactions: alignment × network and alignment × Condition. Neither interaction reached significance for any outcome (all F-test *p*s > .15), and the main-effects model was retained in all cases. In the main-effects models, neural alignment did not significantly predict any of the three outcomes: closeness (*b* = 0.032, SE = 0.058, *t*(388) = 0.544, *p* = .587), enjoyment (*b* = 0.004, SE = 0.064, *t*(388) = 0.059, *p* = .953), or perceived similarity (*b* = 0.074, SE = 0.046, *t*(388) = 1.598, *p* = .111). Taken together, these results again replicate the findings from the main text, demonstrating the reliability of these results.

*Neural exploration predicts conversation outcomes, controlling for model fit.* Here, we repeated the same analysis approach as above to test whether neural exploration predicts conversation outcomes. Neither interaction reached significance for closeness (network: p = .939; Condition: p = .449) or enjoyment (network: p = .993; Condition: p = .261), and the main-effects models were retained for both. Neural exploration did not significantly predict closeness (b = 0.077, SE = 0.079, t(388) = 0.968, p = .334) or enjoyment (b = 0.130, SE = 0.087, t(388) = 1.490, p = .137). For perceived similarity, the exploration × Condition interaction was significant (F(1, 387) = 5.288, p = .022), and the interaction model was retained. Simple slopes revealed that neural exploration significantly predicted greater perceived similarity in Stranger dyads (b = 0.240, SE = 0.073, t(387) = 3.286, p = .001) but not in Friend dyads (b = 0.096, SE = 0.068, t(387) = 1.415, p = .158). The interaction term was negative (b = −0.145, SE = 0.063, t(387) = −2.300, p = .022), indicating that the positive relationship between neural exploration and perceived similarity was significantly attenuated in Friend relative to Stranger dyads. These results mostly match the findings from the main text, again indicating the robustness of our results.
